# *Chlamydomonas* sp. UWO241 exhibits constitutively high cyclic electron flow and rewired metabolism under high salinity

**DOI:** 10.1101/813824

**Authors:** Isha Kalra, Xin Wang, Marina Cvetkovska, Jooyeon Jeong, William McHargue, Ru Zhang, Norman Hüner, Joshua S. Yuan, Rachael Morgan-Kiss

## Abstract

The Antarctic green alga *Chlamydomonas* sp. UWO241 (UWO241) was isolated from the deep photic zone of a permanently Antarctic ice-covered lake. Adaptation to permanent low temperatures, hypersalinity, and extreme shade has resulted in survival strategies in this halotolerant psychrophile. One of the most striking phenotypes of UWO241 is an altered photosystem I (PSI) organization and constitutive PSI cyclic electron flow (CEF). To date, little attention has been paid to CEF during long-term stress acclimation and the consequences of sustained CEF in UWO241 are not known. In this study, we combined photobiology, proteomics, and metabolomics to understand the underlying role of sustained CEF in high salinity stress acclimation. High salt-grown UWO241 exhibited increased thylakoid proton motive flux and an increased capacity for NPQ. A Bestrophin-like Cl^-^ channel was identified in the whole cell proteomes and transcriptome of UWO241 which likely supports ion homeostasis during high transthylakoid pH. Under high salt, a significant proportion of the upregulated enzymes were associated with the Calvin Benson Bassham Cycle (CBB), secondary metabolite biosynthesis, and protein translation. Two key enzymes of the Shikimate pathway, DAHP synthase and chorismate synthase, were also upregulated, as well as indole-3-glycerol phosphate synthase, an enzyme involved in biosynthesis of L-tryptophan and indole acetic acid. In addition, several compatible solutes (glycerol, proline and sucrose) accumulated to high levels in high salt-grown UWO241 cultures. We suggest that UWO241 maintains constitutively high CEF with associated PSI-cytb_6_f supercomplex to support robust growth and strong photosynthetic capacity under a constant growth regime of low temperatures and high salinity.

## INTRODUCTION

During photosynthesis light is transduced into stored energy through two major pathways, linear electron flow (LEF) and cyclic electron flow (CEF). LEF involves the flow of electrons from Photosystem II (PSII) to Photosystem I (PSI) resulting in the production of both ATP and NADPH, which are consumed during carbon fixation in the Calvin Benson Bassham cycle (CBB). CBB requires 3 molecules of ATP and 2 molecules NADPH to fix 1 molecule of CO_2_; however, LEF produces an ATP:NADPH ratio of only 2.57:2 (Kramer and Evans, 2011). CEF constitutes electron transfer from PSI to soluble mobile carriers and back to PSI via cytochrome b_6_f (cyt b_6_f) and plastocyanin (Cardol et al., 2011; Nawrocki et al., 2019). As electrons are shuttled between PSI and cyt b_6_f, a proton gradient is produced that leads to ATP production only. In addition to satisfying the ATP shortage for efficient carbon fixation, CEF-generated ATP may be used for other energy-requiring processes, such as the CO_2_-concentrating mechanism of C_4_ photosynthesis (Takabayashi et al., 2005; Ishikawa et al., 2016), N_2_ fixation in cyanobacteria heterocysts (Magnuson et al., 2011; Magnuson and Cardona, 2016), and survival under environmental stress (Suorsa, 2015).

When the light harvesting antennae absorb light energy in excess of what is required for growth and metabolism, energy homeostasis is disrupted, increasing the risk of formation of reactive oxygen species (ROS) (Hüner et al., 1998). While this phenomenon is associated with excess light, numerous environmental stresses can lead to imbalances in energy demands, including high and low temperatures, high salinity, and nutrient deficiency. Moreover, the duration of the environmental stress can vary over broad time scales, from a few minutes to days or years. Survival of plants and algae requires coordination of short- and long-term acclimatory strategies to maintain energy homeostasis. These acclimation responses are often triggered via the redox status of the plastoquinone pool and initiation of retrograde signaling between the chloroplast and the nucleus (Pfannschmidt, 2003).

CEF is generally accepted as a major pathway utilized under short-term stress by rapidly inducing a transthylakoid pH change and triggering nonphotochemical energy dissipation (Lucker and Kramer, 2013; Yamori et al., 2016b). Early reports linked initiation of CEF with induction of a state transition and formation of a PSI supercomplex (Iwai et al., 2010). Previously, it was assumed that CEF was dependent upon state transitions; however, more recently Takahashi et al. (2013) reported that CEF can initiate in the absence of state transitions. These authors showed that CEF is sensitive to the redox status of intersystem electron transport pool, while Kramer and colleagues showed that hydrogen peroxide acts as a signal for CEF initiation in *Arabidopsis thaliana* (Strand et al., 2015). These previous studies have generally assumed a main role for CEF under short-term, transient stress conditions, while reports of sustained CEF in response to long-term stress are lacking (Lucker and Kramer, 2013).

While the CEF mechanism is not fully understood, formation of protein supercomplexes has been associated with CEF initiation (Minagawa, 2016). The first stable supercomplex was isolated in 2010 by Iwai et al. in *Chlamydomonas reinhardtii*, under short-term exposure to dark/anaerobiosis. This supercomplex is composed of PSI, LHCII, cytochrome b_6_f, PGRL1 and FNR (Ferredoxin NADP Reductase) (Iwai et al., 2010). Takahashi and colleagues (2013) identified another supercomplex in *C. reinhardtii* that is formed under conditions of anoxia and is regulated through the calcium sensing protein, CAS. Recently, the structure of the *C. reinhardtii* PSI supercomplex was solved which showed that dissociation of specific LHCI proteins (Lhca2 and Lhca9) are necessary prior to PSI supercomplex formation (Steinbeck et al., 2018).

Around the globe, there are communities of photosynthetic organisms that have adapted to capture light energy and fix carbon under environmental conditions which are untenable for most model plant and algal species (Dolhi et al., 2013). One example of a prevalent stressful habitat is permanent low temperatures, which encompass the Arctic, Antarctic and alpine environments (Morgan-Kiss et al., 2006). The McMurdo (MCM) Dry Valleys are a large expanse of ice-free land, which forms the largest polar desert on the Antarctic continent. A network of permanently ice-covered lakes provides oases for microbial communities that are vertically stratified through the water column (Priscu et al., 1999; Morgan-Kiss et al., 2006). During the short austral summer, microalgal communities capture light energy and fix carbon, despite numerous permanent environmental stresses, including low temperatures, nutrient deficiency, super-saturated oxygen levels, and hypersalinity (Morgan-Kiss et al., 2006). *Chlamydomonas* sp. UWO241 (UWO241) was isolated from one of the highly studied dry valley lakes (Lake Bonney, east lobe) by J. Priscu and colleagues in the 1990s (Neale and Priscu, 1990; 1995). In its native environment, UWO241 is exposed to year-round low temperatures (0°– 5°C), hypersalinity (700 mM NaCl), and extreme shade (<20 μmol photons m^-2^ s^-1^) of a narrow spectral range (350 – 450 nm).

Early studies on UWO241 focused on growth physiology and its photosynthetic apparatus (Morgan et al., 1998). Most notably, UWO241 appeared to exhibit permanent downregulation of PSI, estimated by a weak P700 photooxidation and an absence of a discernable PSI Chl a low temperature (77K) fluorescence emission peak under a range of treatments (Morgan-Kiss et al., 2002a; 2005; Szyszka et al., 2007; Cook et al., 2019). Earlier reports also suggested that UWO241 appeared to have lost the ability to phosphorylate LHCII and undergo state transitions (Morgan-Kiss et al., 2002b). More recently Szyszka-Mroz et al. (2019) used 33P-labelling to show that UWO241 exhibits some LHCII phosphorylation which is distinct from that of *C. reinhardtii*. The authors also discovered cold adapted forms of the thylakoid protein kinases, STT7 and Stl1, in the psychrophile. Last, they suggested that UWO241 may rely on a constitutive capacity for energy spillover rather than inducible state transitions to regulate energy distribution between PSI and PSII (Szyska-Mroz et al., 2019).

Recent studies have reported that UWO241 maintains sustained CEF under steady-state growth conditions (Szyszka-Mroz et al., 2015; Cook et al., 2019). Constitutively high CEF may represent an adaptive strategy in UWO241 to survive permanent environmental stress, such as low temperatures and high salinity. Hüner and colleagues (Szyszka-Mroz et al., 2015) demonstrated that during growth under high salinity (700 mM NaCl), UWO241 forms a stable PSI supercomplex. The supercomplex was only detectable in cultures acclimated to high salinity, and its stability was disrupted in the presence of the kinase inhibitor, stauroporine.

Why does UWO241 maintain high rates of CEF? On a transitory basis, it is known that CEF is used to satisfy short-term energy needs or to provide pmf to protect PSII by rapid downregulation and induction of qE. However, it is unknown if the consequences of CEF described thus far apply to an organism which appears to utilize CEF on a long-term basis to survive permanent stress. Here, we investigated whether CEF is used for photoprotection or energy generations in UWO241, and also examined the effects of sustained CEF on downstream carbon metabolism.

## RESULTS

### UWO241 possesses constitutively high rates of CEF

UWO241 was isolated from the deep photic zone (17 m sampling depth) of the hypersaline, perennially ice-covered lake (Lake Bonney, McMurdo Dry Valleys, Victoria Land) (Neale and Priscu, 1990; Neale and Priscu, 1995). As a consequence of more than two decades of study, this photopsychrophile has emerged as a model for photosynthetic adaptation to permanent low temperatures (Morgan-Kiss et al., 2006; Dolhi et al., 2013; Cvetkovska et al., 2017). In addition to psychrophily, UWO241 exhibits robust growth and photosynthetic performance under high salt (0.7 M NaCl, Supplemental Fig. S1; Morgan et al., 1998; Pocock et al., 2011).

Low temperature fluorescence spectra of mid-log phase cultures of *C. reinhardtii* and UWO241 grown in control, low salt (LS) growth medium (standard BBM medium, 0.43 mM NaCl) under optimal growth temperatures (20°C and 8°C, respectively) and light conditions (100 μmol m^-2^s^-1^ for both algae) confirmed that *C. reinhardtii* possesses a typical 77K fluorescence emission spectrum with prominent peaks at 685 nm (F_685_) and 715 nm (F_715_), representing LHCII-PSII and PSI, respectively. In agreement with past reports (Morgan et al., 1998; Szyszka et al., 2007), PSI fluorescence was significantly reduced (1.60-fold) in UWO241 relative to *C. reinhardtii* grown under optimal temperature/light conditions in the LS growth medium (Fig. 1A). Moreover, PSI fluorescence was reduced by an additional 1.59-fold in cultures of UWO241 grown in high salinity (HS) growth medium (0.7 M NaCl), relative to LS-grown cells (Fig. 1A).

**Figure 1.**
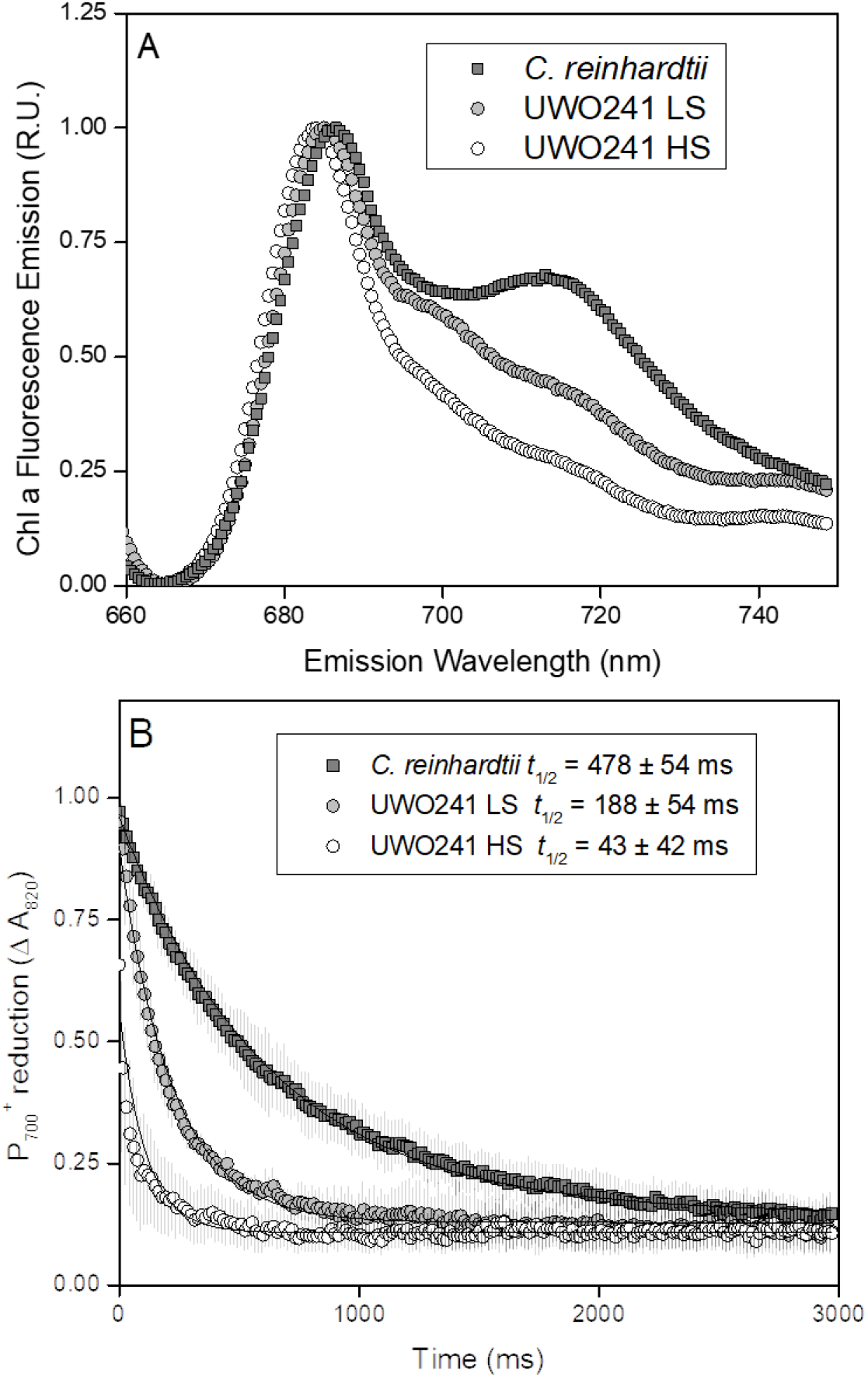
Properties of photosystem I in *C.* sp. UWO241 compared to the model mesophile *C.reinhardtii*. Low temperature (77K) chlorophyll a fluorescence emission spectra of whole cells of *C. reinhardtii* and C. sp. UWO241 (A). P700 re-reduction kinetics. *C. reinhardtii* was grown in low salt BBM medium at 20°C/100 μmol m^-2^s^-1^ (B). UWO241 was grown in either BBM (low salt, LS) or BBM+700 mM NaCl (High salt, HS), and 8°C/100 μmol m^-2^s-^1^. t_1/2_, half-time for P700 re-reduction.

PSI activity was monitored by far red (FR) light inducible P700 photooxidation (Fig. 1B). Following a rise in absorbance at 820 (A_820_), reflecting FR-induced P700 oxidation, we compared rates of P700 re-reduction in the dark in LS cultures of *C. reinhardtii* as well as LS- and HS-grown cells of UWO241 (Fig. 1B). Since FR preferentially excites PSI and not PSII, reduction of P700 following FR exposure is mainly due to alternative electron donors (Ivanov et al., 1998). In agreement with other reports (Morgan-Kiss et al., 2002b; Cook et al., 2019), UWO241 grown in standard LS growth medium exhibited a significantly shorter re-reduction time for P700+ (*t½^red^*) compared with LS-grown *C. reinhardtii* (Fig. 1B). Moreover, HS-grown UWO241 exhibited a 4.4-fold faster *t½^red^* compared with LS-grown cultures (43±42 vs. 188±52 ms respectively; Fig. 1B). These data indicate that relative to the model *C. reinhardtii*, UWO241 exhibits a high capacity for PSI-driven CEF, which is further enhanced during acclimation to long-term high salinity stress.

Higher rates of CEF in HS-grown UWO241 were also confirmed by electrochromic shift (ECS) kinetics which estimates transthylakoid proton flux driven by light-dependent photosynthesis (Fig. 2, Supplemental Fig. S2). The ECS signal was measured by the change in absorbance of thylakoid pigments at 520 nm during application of light dark interval (Baker et al., 2007). The total amplitude of ECS signal (ECS_t_),, was used to estimate the total proton motive force (pmf) across thylakoid membranes (Kramer et al., 2003). UWO241 grown in HS exhibited 6 to 7.5 fold higher ECS_t_ than that of LS-grown cells under all light intensities (Fig. 2A), suggesting HS-grown cells generate higher pmf than LS-grown cells at the same light intensity. High pmf can be caused by either increased proton flux from LEF or CEF, reduced proton efflux, or decreased ATP synthase activity (Kanazawa and Kramer, 2002; Livingston et al., 2010; Carrillo et al., 2016). To verify which process(es) were contributing to high pmf in HS-grown cells, proton conductance (g_H+_) and fluxes through ATP synthase activity (ν_H+_) were analyzed. The inverse of the lifetime of the rapid decay of ECS (g_H+_) represents proton permeability or conductivity of the thylakoid membrane and is largely dependent on the activity of ATP synthesis (Supplemental Fig. S2D; Baker et al., 2007). The g_H+_ of HS-grown cells was ∼50 to 60% of that of LS-grown cells (Figure 2B); however, the proton flux rate (ν_H+_) showed that the amount of ATP produced was still higher in HS-grown cells (Fig. 2C). The relationship between ν_H+_ and LEF can be used to estimate proton contribution from CEF (Baker et al., 2007). In the linear plots of ν_H+_ versus LEF, the slope of the HS-grown cells was higher than that of LS-grown cells (Fig. 2D), indicating that CEF contributes significantly to the total proton exffluxes (ν_H+_) in HS-grown cells. In close agreement with our P700 findings, UWO241-HS exhibited higher rates of CEF compared to UWO241-LS (Figs 1B and 2D; 4.33- and 4.5-fold higher in HS-UWO241 based on P700 and ECS measurements, respectively). Last, HS-grown cells exhibited downregulation of PSII and increased capacity for NPQ (Supplemental Fig. S2A and B), while PSI photochemical yield (Y[PSI]) was higher HS- vs. LS-grown cells due to reduced PSI acceptor side limitation (Y[NA]) (Supplemental Fig. S3).

**Figure 2.**
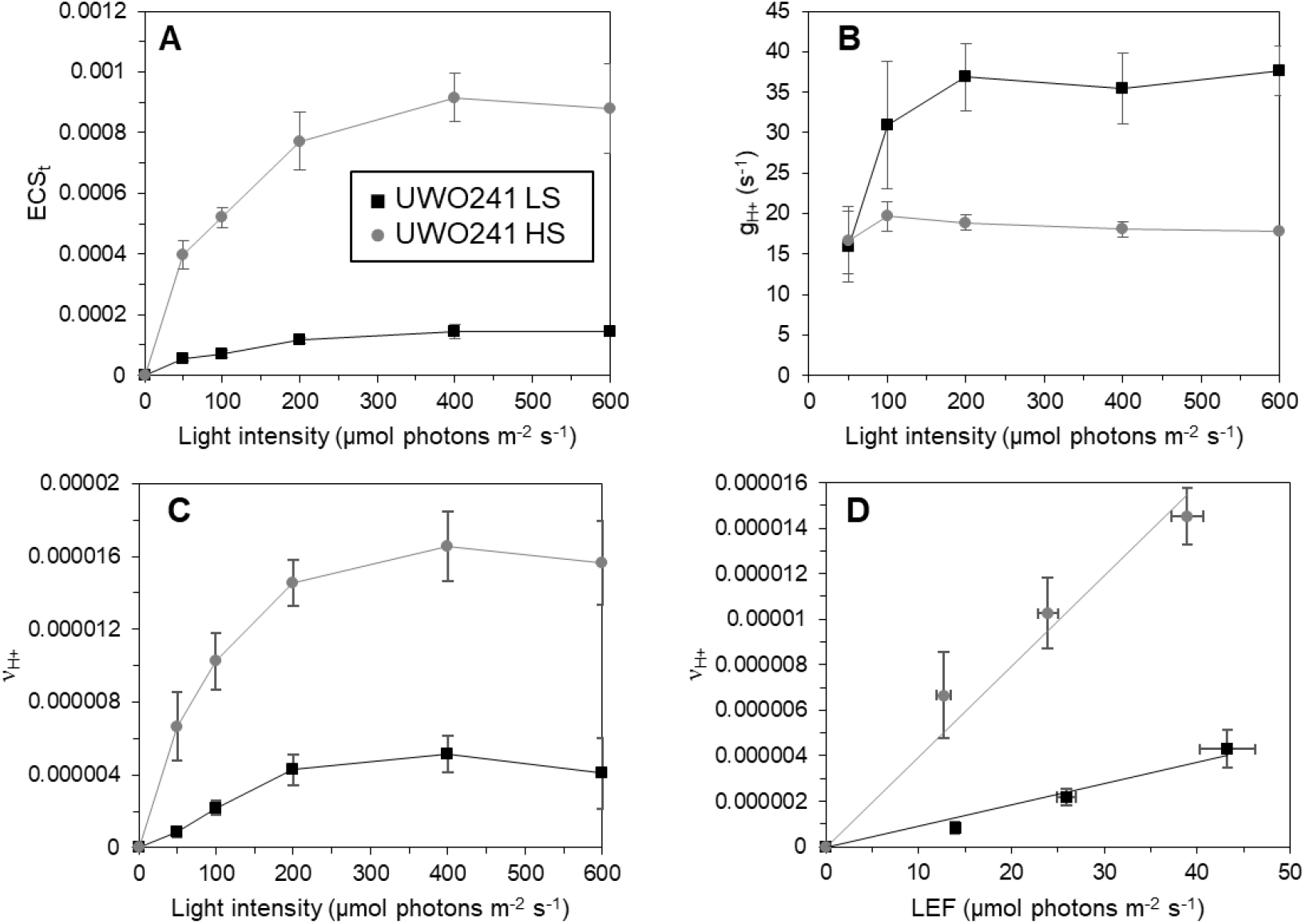
Photosynthetic properties of *C.* sp. UWO241 under various light intensities using Electrochomic shift. The ECS_t_ (total amplitude of ECS, proportional to proton motive force, A), g_H+_ (proton conductance, reflecting the ATP synthesis activities, B) and v_H+_ (proton flux rates, C) of *C*. sp. UWO241 grown in low-salt (black, squares) and high-salt (grey, circles) were measured from dark interval relaxation kinetics (DIRK). The relationship between v_H+_ and LEF (measured by chlorophyll fluorescence, see supplemental Figure S2) were assessed in *C.* sp. UWO241 (D).

### Isolation of a PSI-supercomplex in UWO241

Formation of PSI-supercomplexes have been shown to be essential for induction of CEF in plants and algae (DalCorso et al., 2008; Iwai et al., 2010). An earlier report showed that high salinity-acclimated cultures of UWO241 form a PSI supercomplex (UWO241-SC); however, the yield of the UWO241-SC from fractionated thylakoids was relatively low and only a few proteins were identified (Szyszka-Mroz et al., 2015). In agreement with this report, the sucrose gradient from thylakoids isolated from LS-UWO241 had 3 distinct bands corresponding to major LHCII (Band 1), PSII core complex (Band 2) and PSI-LHCI (Fig. 3A). In contrast, UWO241-HS thylakoids lacked a distinct PSI-LHCI band, but exhibited several heavier bands, including the UWO241-SC band (Band 4; Fig. 3B). We significantly improved recovery of the UWO241-SC by solubilizing thylakoids with the detergent α-DDM rather than β-DDM, which was used by other groups (Fig. 3B). Formation of band 4 of *C. reinhardtii* thylakoids isolated from State 2 conditions, was more diffuse compared to the UWO241-SC (Fig. 3D).

**Figure 3.**
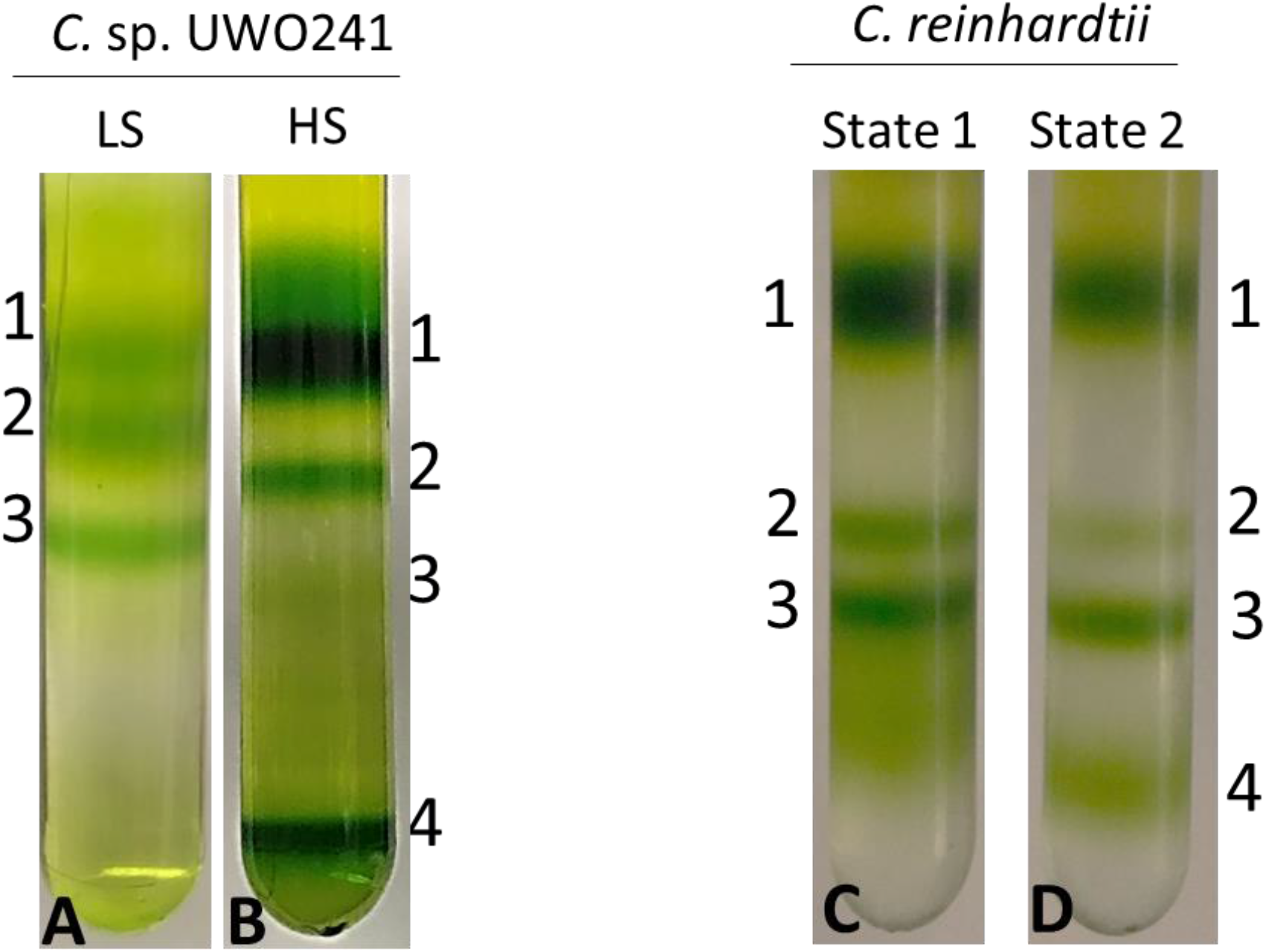
Supercomplex formation in *C.* sp. UWO241 vs. *C. reinhardtii*. Fractionation of major thylakoid chlorophyll-protein complexes from *C.* sp. UWO241 by sucrose density ultracentrifugation from low salt (LS) - and high salt (HS)- grown cultures (A, B). Fractionation of major thylakoid chlorophyll-protein complexes from *C. reinhardtii* exposed to State 1 and State 2 conditions (C, D). Cultures of *C. reinhardtii* were grown under control conditions (20°C/100 umol) and either dark adapted for 10 minutes (State 1, C) or incubated under anaerobic conditions for 30 minutes (State 2, D).

Low-temperature fluorescence spectra were analyzed for the four bands extracted from the sucrose density gradients shown in Fig. 3B and D (i.e. HS-UWO241 and State 2 *C. reinhardtii*, respectively). In *C. reinhardtii*, Band 1 exhibited a major emission peak at 680 nm (Fig. 4C), corresponding to fluorescence from LHCII (Krause and Weis, 1991). Band 2 exhibited emission peak at 685 nm, consistent with PSII core (Fig. 4A). Band 3 exhibited a peak at 685 nm and a strong peak at 715 nm; the latter consistent with PSI-LHCI (Fig. 4A). However, Band 4 exhibited a strong fluorescence peak at 680 nm and a minor peak at 715 nm (Fig. 4A).

**Figure 4.**
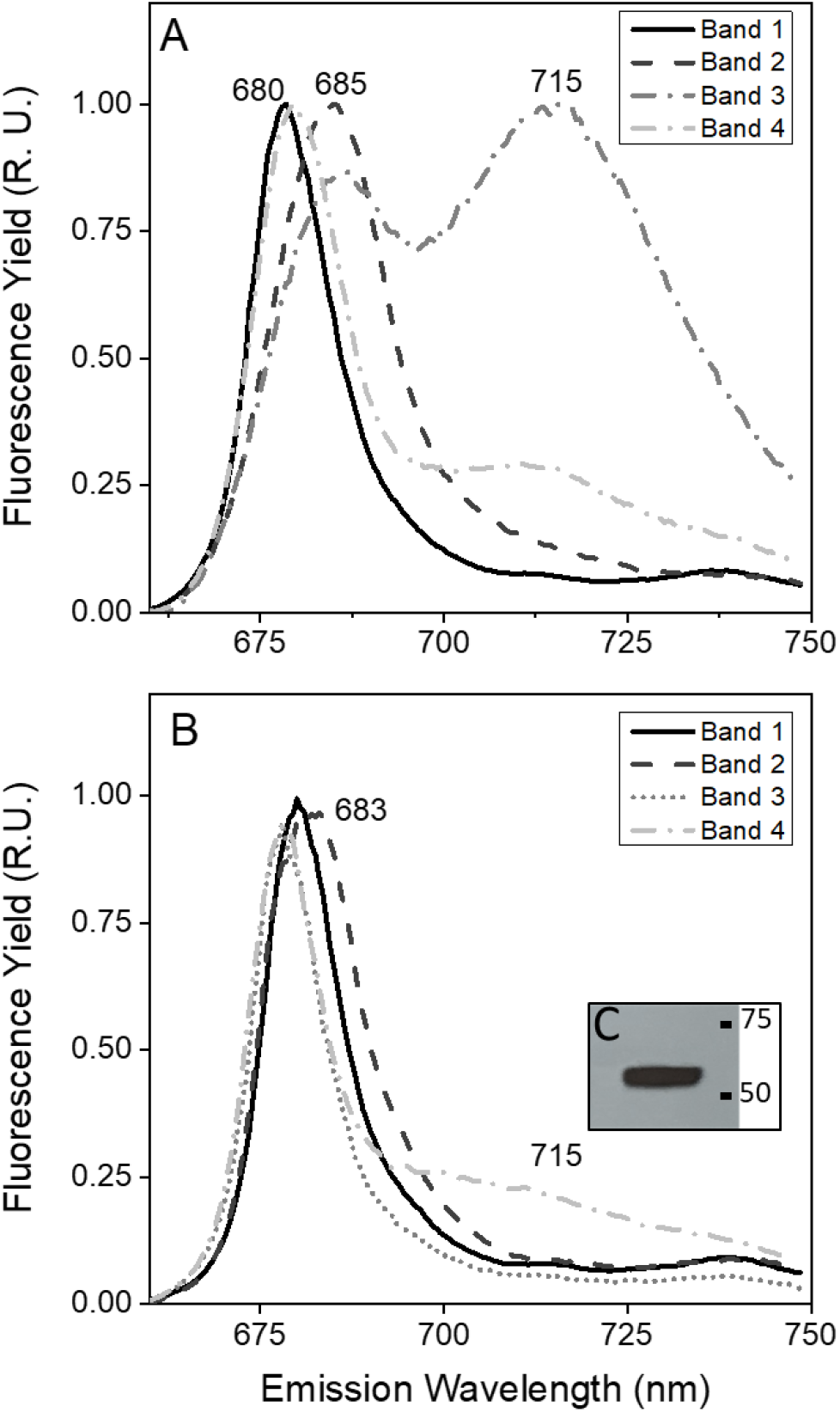
77K fluorescence emission spectra of Chl-protein complexes from *C.reinhardtii* and *C.* sp. UWO241. Low temperature Chl a fluorescence emission spectra of pigment-protein bands isolated from sucrose density gradients shown in Figure 3. Low temperature Chl a emission spectra of bands from *C. reinhardtii* State II conditions (A). Emission spectra of bands from high salt cultures of *C.* sp. UWO241 (B). Immunoblot of the UWO241-SC band with PsaA antibody (C). Spectra represent normalized data and the average of 3 scans.

Fractionated thylakoids from HS-grown UWO241 exhibited emission spectra for Band 1 and Band 2 which were comparable with that from *C. reinhardtii* (Fig. 4B). In contrast, both Band 3 and Band 4 (PSI and SC bands, respectively) exhibited highly reduced or a lack fluorescence associated with PSI. However, we confirmed the presence of the PSI reaction center protein, PsaA, in the UWO241-SC by immunoblotting (Fig. 4C).

### Protein composition of the supercomplex

Protein components of the UWO241-SC were analyzed using LC-MS/MS. We identified a total of 39 proteins in the isolated band 4, significantly more proteins than the previously reported supercomplex of UWO241 isolated using β-DDM (Szyszka-Mroz et al., 2015). The most abundant proteins in the supercomplex were proteins of the PSI reaction center and cytochrome b_6_f. In total we identified seven out of 13 subunits of PSI reaction center (Table 1; Supplemental Table S1). Only two LHCI subunits, Lhca3 and Lhca5, and one LHCII minor subunit, CP29 were associated with the UWO241-SC. The Calcium sensing receptor (CAS) was identified as the third most abundant protein in the UWO241-SC. We also identified four subunits of ATP synthase in the UWO241-SC (α, β, γ, δ). In agreement with an earlier report, we found FtsH and PsbP in the UWO241-SC band (Szyszka-Mroz et al., 2015).

**Table 1.** The UWO241 supercomplex contains subunits of PSI, Cytb_6_f, ATP synthase, as well as several antenna proteins. Proteomic analysis of Band 4 isolated from cultures acclimated to high salinity (700 mM NaCl). NSAF (percent normalized spectral abundance factor) for selected proteins are shown.

| Protein | NSAF (%) | UniProtKB | Organism |
| --- | --- | --- | --- |
| **PSI** |  |  |  |
| **PsaD** | 4.53 | Q39615 | *C. reinhardtii* |
| **PsaE** | 4.24 | P12356 | *C. reinhardtii* |
| **PsaF** | 3.12 | P12356 | *C. reinhardtii* |
| **PsaH** | 1.84 | P13352 | *C. reinhardtii* |
| **PsaK** | 6.13 | P14225 | *C. reinhardtii* |
| **PsaL** | 2.43 | Q39654 | *C. sativus* |
| **LHCI & LHCII Antenna** |  |  |  |
| **Lhca3** | 1.35 | Q9SY97 | *A. thaliana* |
| **Lhca5** | 2.82 | Q9C639 | *A. thaliana* |
| **CP29** | 2.25 | Q93WD2 | *C. reinhardtii* |
| **LHCII type I** | 2.22 | P20866 | *P. patens* |
| **CB2** | 9.46 | P14273 | *C. reinhardtii* |
| **Cyt b<sub>6</sub>f** |  |  |  |
| **PetA** | 5.35 | P23577 | *C. reinhardtii* |
| **PetB** | 2.33 | Q00471 | *C. reinhardtii* |
| **PetC** | 1.15 | P49728 | *C. reinhardtii* |
| **ATP Synthase** |  |  |  |
| **AtpA** | 2.45 | P26526 | *C. reinhardtii* |
| **AtpB** | 2.07 | P06541 | *C. reinhardtii* |
| **Other** |  |  |  |
| **CAS** | 4.65 | Q9FN48 | *A. thaliana* |
| **FtsH1** | 0.49 | Q5Z974 | *O. sativa* |
| **FtsH2** | 1.26 | O80860 | *A. thaliana* |
| **PsbP** | 2.52 | P11471 | *C. reinhardtii* |

Bands 3 and 4 contained many PSI core proteins but lacked almost all Lhca proteins (Supplemental Table S1). These results agreed with an earlier report that failed to detect most Lhca proteins in UWO241 thylakoids (Morgan et al., 1998). To determine whether the absence of Lhcas in the UWO241 proteome was due to a loss of Lhca genes, we searched a UWO241 transcriptome generated from a culture grown under low temperature/high salinity (Raymond et al., 2009). Surprisingly, we identified 9 Lhca homologues which were transcribed under high salinity, suggesting that all or most of the LHCI genes are expressed in UWO241 (Supplemental Fig. S4).

### Expression of a Bestrophin-like transporter

A recent report detected expression of a bestrophin-like anion channel protein in whole cell proteomes of UWO241 (Cook et al., 2019). A search of the UWO241 transcriptome revealed two homologues that were related to other algae and plant BEST1 proteins (Fig. 5A). Both UWO241 BEST1 homologues (contig 864 and 973) contain four putative transmembrane domains (Fig. 5B and C). Contig 864 possesses a putative chloroplast transit peptide and cleavage site, suggesting that it is localized to the chloroplast. Modeling of the UWO241 BEST1 protein using KpBest as a template, indicated that the UWO241 BEST channel may form a pentamer, with the Cl^-^ entryway and exit is located on the stromal and luminal sides, respectively of the thylakoid membrane (Fig. 5D), similar to a recent study on AtBEST1 (Duan et al., 2016).

**Figure 5.**
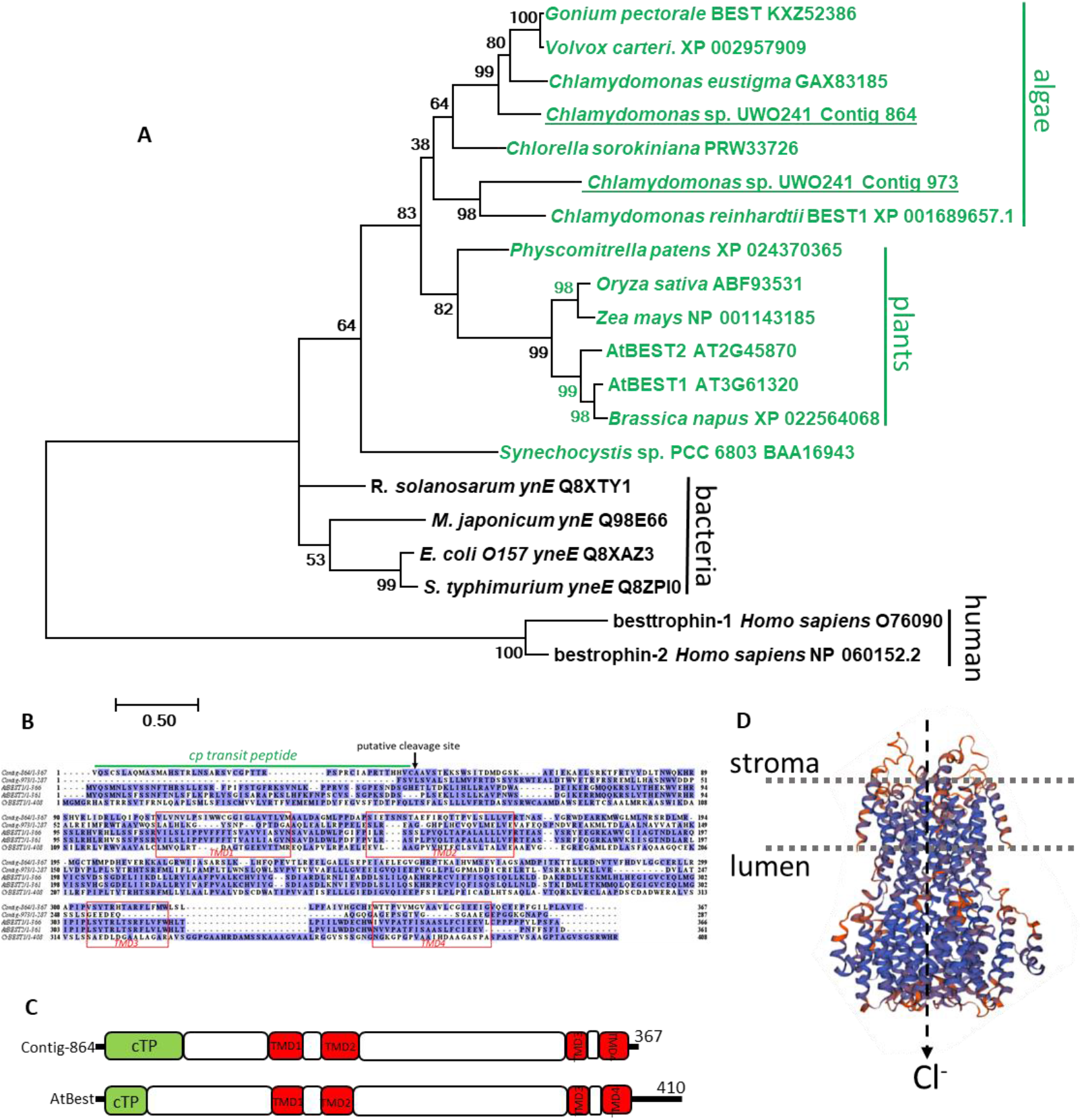
Expression of a Bestrophin-like transporter in UWO241. Maximum likelihood tree of the BEST1 homologue identified in the UWO241 transcriptome from HS-grown cultures (A). Numbers are bootstrap values from 500 replicate trees. Protein alignment of BEST proteins (B). Protein schematics for putative C. sp. UWO241 BEST protein (contig 367) and *A. thaliana* Best1 (C) (Duan et al., 2016). Green and orange blocks indicate putative chloroplastic transit peptides and trans-membrane domains, respectively. Predicted tertiary structure of UWO241 BEST channel generated by SwissModel (https://swissmodel.expasy.org/) using *Klebsiella pneumoniae* Best complex as a template (D).

### Whole cell proteome analysis

We wondered whether constitutively high CEF could be linked to changes in downstream metabolism in UWO241. To address this question, whole-cell proteomes extracted from cultures of UWO241-HS and -LS grown under optimal temperature/light conditions were compared. Overall 98 proteins from various functional categories were identified as significantly affected in the two treatments, out of which 46 were upregulated and 62 downregulated in HS-acclimated cells. Proteins associated with photosynthesis (18%) and translational machinery (18%) were most affected in the treatment followed by primary and secondary metabolism (16%) (Supplemental Fig. S5).

#### Photosynthesis

Differentially regulated proteins participating in photosynthesis were identified in UWO241-HS. Three photosystem reaction center proteins, PsaB and PsaN of PSI, and D2 protein of PSII, as well as extrinsic proteins of the water oxidizing complex, PsbO and PsbQ, were downregulated in UWO241-HS (Figs. 6 and 7A; Supplemental Table S2). One protein of chloroplastic ATP synthase, the epsilon subunit, was upregulated (3.8 fold) in UWO241-HS. Last, both FtsH proteins that were detected in the UWO241-SC, FtsH1 and FtsH2, were upregulated in UWO241-HS (Table 1; Supplemental Table S2).

**Figure 6.**
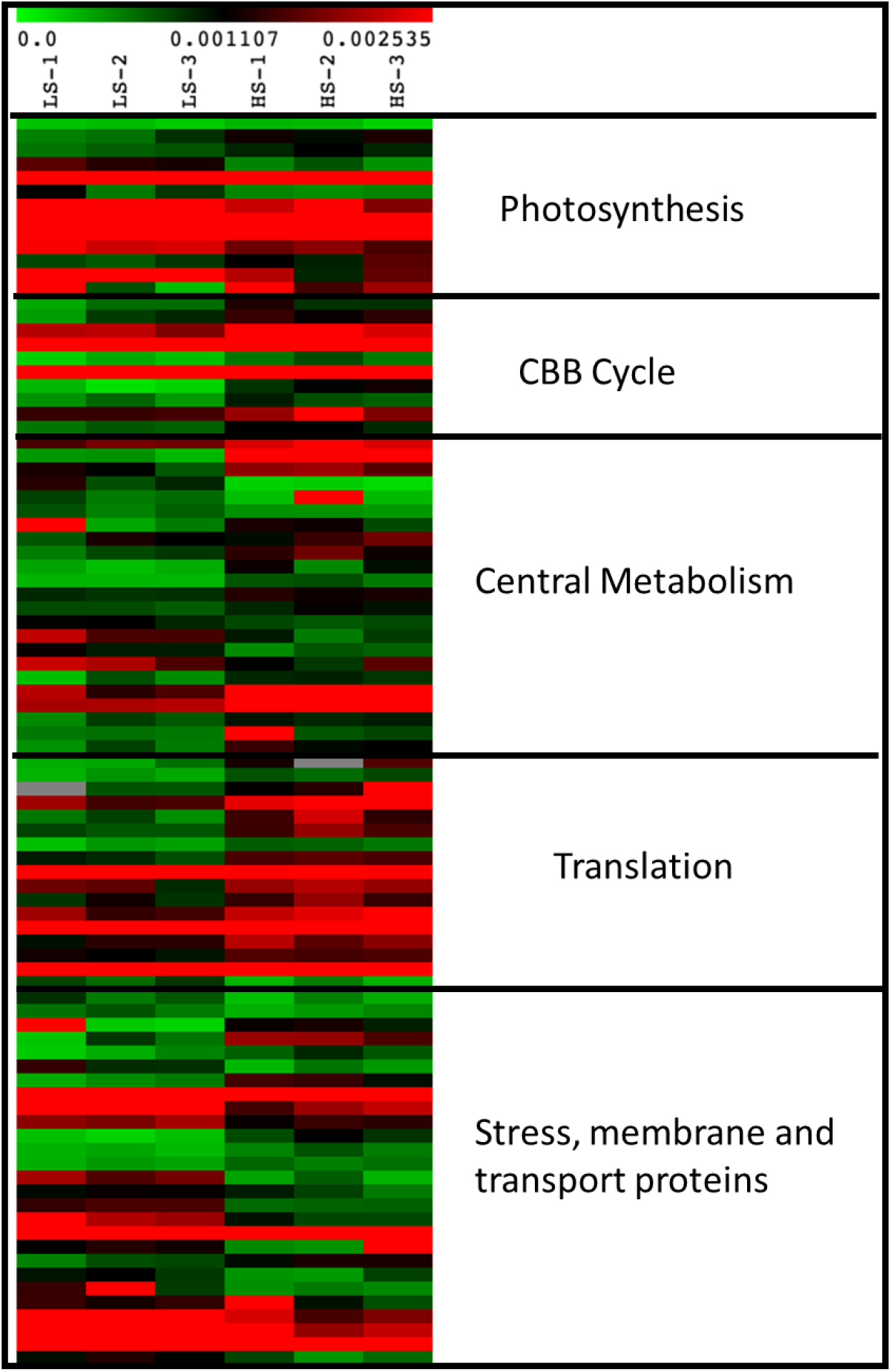
Heat map of differentially regulated proteins in UWO241 under low (LS) and high salinity (HS) conditions. The normalized spectral abundance factor (NSAF) values are plotted for each replicate in the two conditions (n=3) using color based approach (green: low abundance, red: high abundance). The proteins are categorized into broad processes they belong to.

**Figure 7.**
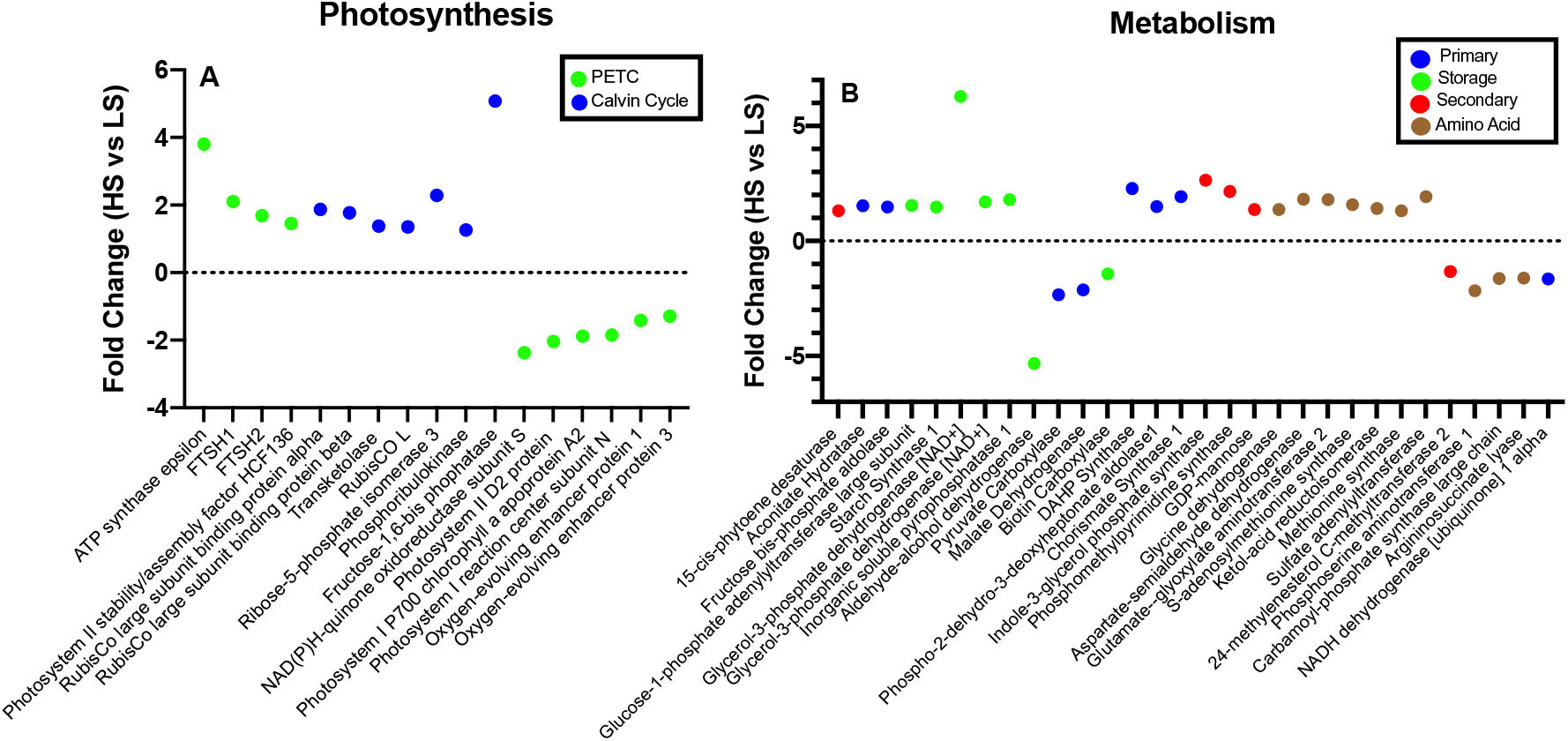
Changes in photosynthetic machinery and downstream metabolism in UWO241 associated with high salinity. Proteins participating in photosynthetic machinery (PETC: green, Calvin cycle: Blue) that are significantly affected in UWO241 under high salinity (A). Proteins participating in downstream metabolism that belong to primary (blue), storage (green), secondary (red) and amino acid biosynthesis (brown) metabolic pathways and are significantly affected in UWO241 under high salinity are shown (B). Proteins with fold change > 1.2 are shown (n=3). PETC: Photosynthetic electron transport chain.

Several proteins associated with the CBB were significantly upregulated in UWO241-HS cells (Figs. 6 and 7A; Supplemental Table S2). The RubisCO large subunit was upregulated in UWO241-HS, along with two chaperone proteins involved in RubisCO assembly, RuBA and RuBB. A class 2 fructose-1,6 –bisphosphatase (FBPase) was the third highest upregulated protein (5-fold), and fructose bisphosphate aldolase and transketolase were also upregulated in UWO241-HS (Fig. 7A).

#### Metabolism

The TCA cycle protein aconitate hydratase was upregulated in UWO241-HS, while pyruvate carboxylase, which is involved in anaplerotic reactions, was downregulated (Fig. 7B; Supplemental Tables S2 and S3). In addition, malate dehydrogenase (MDH) was downregulated in UWO241-HS. MDH participates in the malate shunt and helps shuttle excess reducing power from the chloroplast to the mitochondria by converting oxaloacetate to malate. In this process excess NADPH are used and NADP pool is regenerated (Scheibe, 2004).

Lipids and starch are major forms of stored energy in green algae. Fructose bis-phosphate aldolase, involved in gluconeogenesis and feeding into starch synthesis, as well as glucose-1-phosphate adenylyltransferase and starch synthase 1 (the last enzyme in starch synthesis pathway) were upregulated in UWO241-HS (Fig. 7B; Supplemental Table S2). In addition, the enzyme glycerol-3-phosphate dehydrogenase (G3PDH), involved in glycerol biosynthesis, was the highest upregulated enzyme in UWO241-HS cultures (6-fold) (Fig. 7B; Supplemental Table S2). G3PDH is involved in conversion of DHAP to sn-glycerol-3-phosphate that leads to glycerol production through glycerol kinase (GK) or G3P phosphatase (GPP) (Driver et al., 2017). The G3P produced in this reaction is also a precursor for TAG synthesis and can also lead to increased lipid production under salinity stress (Herrera-Valencia et al., 2012). On the other hand, Alcohol-aldehyde dehydrogenase (AADH) was significantly downregulated in UWO241-HS (5-fold; Fig. 7B; Supplemental Table S3).

Secondary metabolic pathways act as large energy sinks and help organisms deal with stress by producing useful secondary metabolites (Darko et al., 2014). Two key enzymes from the Shikimate pathway were significantly upregulated under high salinity in UWO241: (i) the first enzyme in the pathway, DAHP (3-Deoxy-D-arabinoheptulosonate 7-phosphate) synthase (2.3-fold), and (ii) the last enzyme in the pathway, chorismate synthase (1.9-fold) (Fig. 7B; Supplemental Table S2). Chorismate, the product of the Shikimate pathway, is a substrate for both aromatic amino acids and many phenylpropanoid secondary metabolites. We found that indole-3-glycerol phosphate synthase (IGP synthase) was upregulated significantly in the HS conditions. IGP synthase is a branching enzyme that can either enter tryptophan pathway or lead to *de nov*o biosynthesis of the plant phytohormone indole acetic acid (IAA) (Ouyang et al., 2000).

### Primary metabolome analysis

Comparing the whole cell proteome of LS and HS grown cultures suggested that salinity has a strong effect on primary and secondary metabolism in UWO241. To further explore this, UWO241 metabolic extracts from LS and HS cultures were analyzed using GC-MS. We detected a total of 771 unique metabolic signatures, 179 of which were positively identified based on their mass spectra and retention times according to the FiehnLab mass spectral database (Kind et al., 2009). PCA analysis of all unique metabolites demonstrates that the metabolic status of UWO241 cells grown under HS conditions is significantly different from that grown under LS along PC1, which accounts for most of the variability between samples (53.8%) (Supplemental Fig. S7). Overall, 186 out of 771 metabolites (24%) were identified as being significantly different among the two growth conditions (t-test; p<0.01). A heat map of all measured (identified and unidentified) metabolites showing the relative changes in primary metabolite abundances indicated clustering and a discrete population of metabolites that accumulate at high levels in HS-grown cultures when compared to LS-grown cells (Fig. 8). To better understand the effect of high salinity on the metabolic profile of UWO241, we performed a detailed analysis on the subset of primary metabolites that were positively identified. Overall, 59 metabolites (32%) from different chemical categories were significantly different (p<0.01), out of which 9 were present in higher abundance (Supplemental Table S4) and 50 were present in lower abundance (Supplemental Table S5) in HS-grown UWO241 cultures.

**Figure 8.**
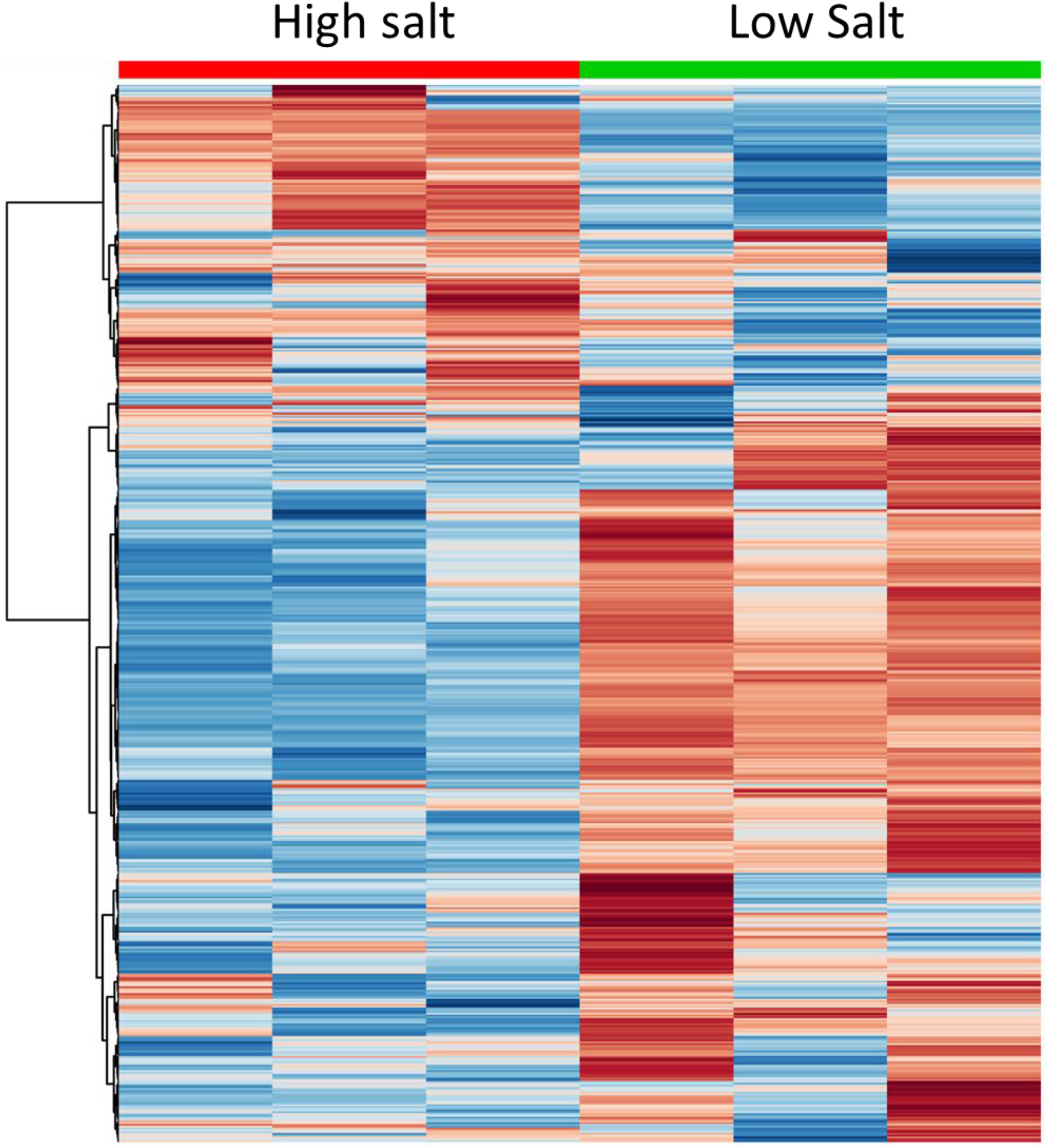
Heat map showing the relative changes in primary metabolite abundances in *C.* sp. UWO241 cultures grown under high or low salt conditions. The analysis includes all quantified metabolites. For each growth condition, three biological replicates are represented using a color-based metabolite profile as indicated (red – high abundance; blue – low abundance). Hierarchical clustering is based on Euclidean distances and Ward’s linkage

#### Metabolites that accumulate in high amounts in HS grown cultures

We observed high levels of glycerol in the primary metabolome of HS-grown UWO241 (8.7 FC), and high accumulation of the compatible solutes, sucrose (18.2 FC) and proline (27.1 FC) (Supplemental Table S4). We also observed a high accumulation of phytol (12.6 FC), suggesting chlorophyll degradation. Tocopherol was also detected in our HS experiment in high amounts (9.5 FC), however its accumulation was variable and thus not statistically different between samples (p>0.05).

#### Metabolites that accumulate in lower amounts in HS grown cultures

UWO241 cultures grown in high salinity exhibited decreased amounts of 17 amino acids and compounds associated with amino acid metabolism (Supplemental Table S5). Most notable metabolites from this class were lysine (29.4 FC) and ornithine (14.0 FC), which could signify a shift in amino acid metabolism to proline during exposure to high salinities. We also observed lower levels of the amino acid tryptophan (2.8 FC) in HS grown cultures. Metabolites involved in purine and pyrimidine metabolism were present in lower amounts in UWO241 exposed to high salinity, suggesting that these cells have shifted their metabolism from maintenance of the cell cycle and nucleic acid synthesis to producing osmoprotectants and compatible solutes. We also observed a reduction of 3-phosphoglycerate (3-PGA; 2.9 FC) in HS-grown cultures.

## DISCUSSION

Our study shows that UWO241 maintains robust growth and photosynthesis under the combined stress of low temperature and high salt. This ability differs markedly from other model plants and algae that typically display downregulation of photosynthesis and growth when exposed to environmental stress, mainly as a consequence of bottlenecks in carbon fixation capacity (Hüner et al., 1998; Hüner et al., 2003; Ensminger et al., 2006; Hüner et al., 2016). Previous research has thoroughly described adaptive strategies for survival under permanent low temperatures, while survival under hypersalinity has received less consideration (Morgan et al., 1998; Morgan-Kiss et al., 2002a; Szyszka et al., 2007; Possmayer et al., 2011).

One of the more distinct photosynthetic characteristics of UWO241 is the presence of a strong capacity for PSI-driven CEF (Morgan-Kiss et al., 2002b; Szyszka-Mroz et al., 2015; Cook et al., 2019). We validated that CEF rates are high in HS-grown cultures using the ECS signal, which was purported to mitigate problems with using P700 absorbance changes for CEF estimates (Lucker and Kramer, 2013). While CEF appears to be essential in plants and algae for balancing the ATP/NADPH ratio and protecting both PSI and PSII from photo-oxidative damage (Munekage et al., 2004; Joliot and Johnson, 2011; Huang et al., 2012), most studies report that CEF is part of short-term stress acclimation. Our work here as well as others suggests a larger role for CEF during long-term adaptation under permanent environmental stress (Morgan-Kiss et al., 2002b; Szyszka-Mroz et al., 2015; Cook et al., 2019a).

Constitutively high rates of CEF in UWO241 are associated with a general reorganization of PSI and PSII complexes (Szyszka-Mroz et al., 2019) and the formation of a Cyt bf-PSI supercomplex (Szyszka-Mroz et al., 2015). In this current study, following optimization of thylakoid protein complex solubilization by substituting β-DDM with α-DDM, the vast majority of PSI shifts from free PSI in the LS-grown cultures to association with the UWO241-SC in the HS-grown cultures. PSI supercomplexes have been described in several plant and algal species (Iwai et al., 2010; Li et al., 2018; Steinbeck et al., 2018). The UWO241-SC is distinct from that of *C. reinhardtii* because: i) its assembly is independent of short-term exposure to dark anaerobic conditions or other state transition-inducing treatments (Fig. 3), ii) the vast majority of PSI in UWO241 is associated with the UWO241-SC (Fig. 3), and iii) isolated UWO241-SC and PSI bands as well as whole cells lack typical PSI fluorescence emission at 77K, despite the presence of several PSI core proteins (Fig. 4; Table 1; Supplemental Table S1; Morgan et al., 1998; Morgan-Kiss et al., 2002a, b; Cook et al., 2019). Thus, CEF combined with significant structural and functional changes to PSI are major targets for long-term stress acclimation in UWO241 (Fig. 9A).

**Figure 9.**
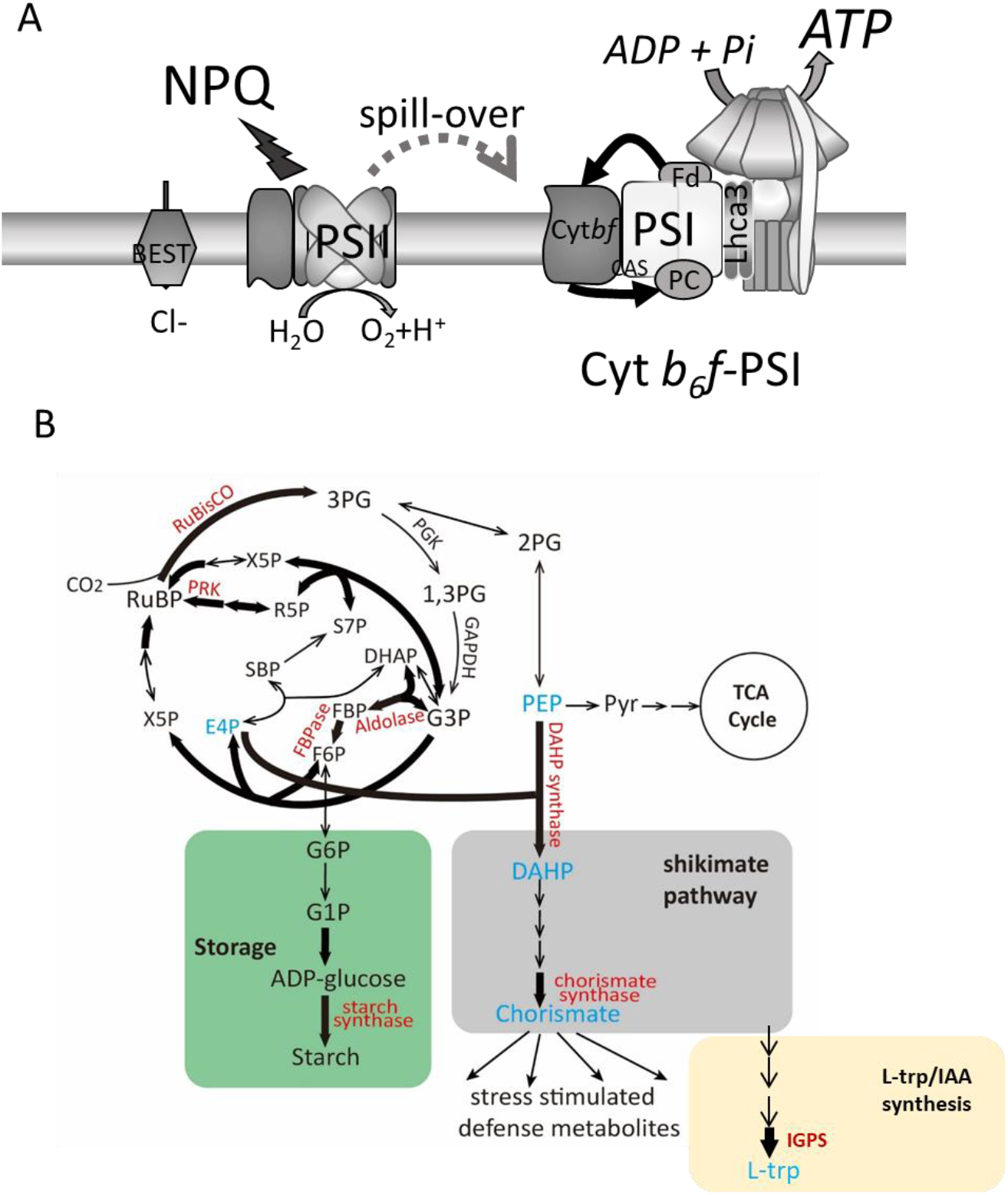
A restructured photosynthetic apparatus supports rewiring of central metabolism in *C.* sp. UWO241. The photosynthetic apparatus of UWO241 is assembled to promoted high rates of CEF which is sustained by formation of a PSI supercomplex (A). CEF supports photoprotection of PSII and PSI and provides additional ATP for downstream metabolism. High ATP is consumed, in part, by the CBB cycle as well as an upregulated shikimate pathway and carbon storage pathways (starch, glycerol) (B). Model is based on data presented here and other studies (Cook et al., 2019; Szyszka-Mroz et al., 2015; 2019)

While the UWO241-SC contains most of the PSI core proteins, both the UWO241-SC and PSI bands, as well as whole cell proteomes isolated from LS and HS conditions lacked homologues for most LHCI proteins (Table 1; Supplemental Table S1). This agrees with an earlier study which was unable to detect most of the LHCI polypeptides by immunoblotting in UWO241 thylakoids (Morgan et al., 1998). Cook et al. (2019) also reported that the absence of LHCI proteins in UWO241 was not associated with adaptation to chronic iron deficiency, an additional stress experienced by natural communities of this alga. Transcriptomic analyses detected nine Lhca homologues in UWO241 grown under low temperature/high salinity (Supplemental Fig. S4). Therefore, it appears that while most of the Lhca genes are encoded for and transcribed, few of the LHCI proteins are produced under the growth conditions tested thus far. These results fit well with numerous unsuccessful attempts to elicit typical 77K PSI long wavelength fluorescence emission in UWO241 (Morgan et al., 1998; Morgan-Kiss et al., 2002a; Morgan-Kiss et al., 2002b; 2005; Szyszka et al., 2007; Cook et al., 2019). It also explains the differences in the 77K emission spectra of the UWO241-SC and PSI bands between UWO241 and *C. reinhardtii* (Fig. 4). Last, a recent study reported that UWO241 transfers light energy from PSII to PSI via constitutive energy spillover through an undescribed mechanism (Szyska-Mroz et al., 2019). Thus, UWO241 favors downregulated LHCI and constitutive energy spill-over in response to its extreme habitat, most likely the natural light environment of extreme shade enriched in blue wavelengths (Neale and Priscu, 1995).

The direct product of CEF is extra transthylakoid proton motive force at the expense of NADPH production (Lucker and Kramer, 2013; Dumas et al., 2016; Yamori et al., 2016a), with the consequences of CEF impacting either cellular energy production or photoprotection. CEF-dependent formation of ΔpH protects PSII by activating the energy-dependent quenching (qE), a major process for dissipation of excess light energy in PSII (Yamori et al., 2016). Alternatively, CEF-generated pmf can be used for production of additional ATP for high-energy consuming processes including protein synthesis, transport processes, ion homeostasis (He et al., 2015), CO_2_ concentrating mechanisms (Horváth et al., 2000; Lucker and Kramer, 2013), or production of secondary metabolites (Murthy et al., 2014). Last, CEF prevents PSI photoinhibition by downregulating LEF and alleviating over-reduction of the acceptor side of PSI, thereby preventing ROS-induced PSI damage (Munekage et al., 2008; Shimakawa et al., 2016; Chaux et al., 2017; Huang et al., 2017). A strong constitutive CEF mechanism in UWO241 could be beneficial for one or most of the above purposes. First, HS-grown cultures possess a higher capacity for NPQ (Supplemental Fig. S2), supporting a role for CEF in constitutive photoprotection ability. Expression of a thylakoid BEST ion channel also suggests CEF may be used for NPQ. High CEF rates also correlate with a higher Y(PSI) and a lower PSI acceptor side limitation in HS-grown cultures (Supplemental Fig. S3), suggesting enhanced PSI photoprotection in UWO241-HS cells.

ATP synthase subunits were associated with the UWO241-SC (Table 1), suggesting CEF contributes extra ATP in UWO241. HS-grown cultures exhibited significantly higher ECS_t_ and ν_H+_ compared to LS-grown cultures, suggesting a high flux of protons through the chloroplastic ATP synthase in spite of slow activity of ATP synthase (compare Fig. 2A and C with Fig. 2B). Slower activity of ATP synthase could be overcome by higher ATP synthase subunits in the HS-grown UWO241 which is reported here (Figs. 6 and 7) and in an earlier report (Morgan et al., 1998). Recently it was shown that in a salt-tolerant soybean, increased CEF contributes to excess ATP that is used to drive import of Na^+^ in the vacuole (He et al., 2015). Taken together, constitutively high rates of CEF in UWO241 are likely to provide dual benefits, that of constitutive photoprotection of both PSI and PSII and extra ATP to support downstream processes important for low temperature and/or high salinity adaptation (Fig. 9A).

Comparison of whole cell proteomes and metabolomes revealed significant shifts in primary and secondary metabolism in LS- and HS-grown UWO241. First, HS-grown cultures have a strong carbon fixation potential. Key enzymes within the CBB cycle are upregulated under HS, including large subunit of RubisCO (LSU, EC 4.1.39) and a chaperone complex involved in RubisCO assembly (RuBA and RuBB proteins). Several enzymes important in regeneration of ribulose-1,5-bisphosphate (RuBP), including fructose-1,6-bisphosphatase (FBPase, EC 3.1.3.11), fructose-bisphophate aldolase (EC 4.1.2.13), transketolase (EC 2.2.1.1), ribose-5 phosphate isomerase (EC 5.2.1.6), and phosphoribulokinase (PRK, EC 2.7.1.19), are also higher in HS-grown cells (Fig. 9B). Overexpression of key bottleneck CBB enzymes such as FBPase and SBPase enhances carbon fixation and RuBP regeneration (Lefebvre et al., 2005; Tamoi et al., 2006), while also supporting improved photosynthesis during stress (Driever et al., 2017). Low levels of 3-PGA, the direct biproduct of RubisCO activity also suggests strong carbon sinks for fixed CO_2_ in HS-grown cultures (Supplemental Table S5). Last, overproduction of these key CBB enzymes is supported by a robust protein translation ability, as several ribosomal proteins are also overexpressed in HS-grown cells (Supplemental Table S2).

Enhanced CBB pathway activity would support robust photosynthetic activity and growth in UWO241. However, proteomic evidence revealed other potential carbon sinks, including carbon storage in the form of starch (Supplemental Table S2). Two key enzymes of starch synthesis were upregulated under high salinity, G1P adenylyltransferase (AGPase; EC 2.7.7.27) and starch synthase (EC 2.4.1.242). AGPase catalyzes the formation of glucose-1-phosphate to ADP-glucose and consumes 1 ATP. ADP-glucose serves as substrate for starch synthase to extend the glucosyl chain in starch. In contrast with these findings, plant and algal fitness and survival under low temperatures or high salinity is associated with starch degradation (reviewed in Thalmann et al., 2016). Under abiotic stress starch is remobilized into sugars and other metabolites to provide carbon and energy when photosynthesis is compromised. In contrast with cold- or salt-sensitive plants and algae, the photosynthetic apparatus of UWO241 is remodeled to support photosynthesis under continuous low temperatures and high salinity (Morgan et al., 1998; Szyszka et al., 2007; Pocock et al., 2011). Thus, accumulation of starch in UWO241 may act as a strong carbon sink to support high rates of carbon fixation (Fig. 9B). Starch stored in the chloroplast is also transitory, and is often rapidly turned over (Thalmann et al., 2016). Starch content was comparable between LS- and HS-grown UWO241 cultures (Supplemental Fig. S6). Thus, transiently stored starch could be an additional adaptive strategy in UWO241, acting as an energy and carbon buffer which can be rapidly mobilized when needed. This theory is supported by other publications that reported accumulation of starch under cold or salinity stress (Siaut et al., 2011; Wang et al., 2013), suggesting that transitory starch synthesis and mobilization may be important during stress acclimation.

Glycerol is a compatible solute that accumulates at molar levels in the salt-tolerant alga *Dunaliella* (Avron, 1986; Brown, 1990; Goyal, 2007a,b). Glycerol is synthesized through two independent pathways localized in the chloroplast and the cytosol. In the presence of light, the chloroplast pathway dominates at the expense of starch synthesis (Gimmler and Möller, 1981), while in the dark, stored starch is degraded to provide substrates for the cytosolic pathway (Ben-Amotz and Avron, 1973). Under high salinity stress, *D. tertiolecta* utilizes both the chloroplast and cytosolic pathways for glycerol synthesis (Goyal, 2007a, b). Overexpression of *D. bardawil* SBPase in *C. reinhardtii* led to increased accumulation of glycerol and improved photosynthesis under salinity stress (Fang et al., 2012). This current study showed that UWO241 also accumulates glycerol in response to increased salinity (Supplemental Table S4). Synthesis of glycerol could occur through either the chloroplast or cytosolic pathway, since isoforms of both the cytosolic and chloroplast glycerol-3 phosphate dehydrogenases (GPDH, EC 1.1.1.8) were upregulated under high salinity (Supplemental Table S2). GPDH is responsible for the first step in glycerol synthesis, conversion of dihydroxyacetone phosphate (DHAP) to glycerol 3-phosphate (G3P), and also supplies G3P for chloroplastic glycerolipid synthesis (Chandra-Shekara et al., 2007). The cytosolic GPDH is highly overexpressed in UWO241, suggesting that glycerol production via starch breakdown may be the dominant pathway in this organism.

The shikimate pathway is an essential link between primary and secondary metabolism for producing precursors for aromatic amino acids (tryptophan, phenylalanine, tyrosine) as well as many other aromatic metabolites such as, indole compounds, alkaloids, lignin and flavonoids (Maeda and Dudareva, 2012). It is a high carbon flux pathway, accounting for approx. 30% of all fixed CO_2_ in an organism (Tohge et al., 2013). Various biotic and abiotic stresses upregulate genes of the Shikimate pathway as well as downstream biosynthesis pathways which use the main product of the Shikimate pathway, chorismate, as a substrate (Tzin and Galili, 2010). Chorismate is an important substrate for production of a number of molecules important for plant defense, including salicylic acid and auxins (Berens et al., 2017). HS-grown UWO241 exhibited upregulation of two key Shikimate enzymes, 3-deoxy-D-arabino-heptulosonate 7-phosphate synthase (DHAP synthase; EC 2.5.1.54) and chorismate synthase (EC 4.2.3.5), catalyzing the first and last enzymatic reactions, respectively (Fig. 9B; Supplemental Table S2). The substrates for DAHP synthase are erythrose-4-phosphate (E4P) and PEP. E4P is a product of the CBB cycle enzyme transketolase. Upregulation of transketolase in HS-grown cells indicates that E4P would be supplied in high levels to support high flux through the Shikimate pathway. On the other hand, supply of PEP for chorismate synthesis is likely to come from glycolysis, as indicated by the higher level expression of G3P dehydrogenase under the HS condition (Supplemental Table S2). Thus, the CBB cycle and glycolysis are likely to be coordinated in order to provide substrates to support high flux through the shikimate pathway. Last, there is evidence linking CEF with the Shikimate pathway. One recent study involving a CEF mutant of *Arabidopsis thaliana* (lacking pgr5 protein) showed that the levels of Shikimate metabolites were significantly reduced in the CEF mutant as compared to wild type, suggesting a link between CEF and chorismate synthesis (Florez-Sarasa et al., 2016).

Acclimation to a variety of stresses in plants and algae often involves upregulation of heat shock proteins, stress metabolites, as well as signaling molecules such as plant hormones and signal transduction pathways (eg: Ca^2+^) (Montero-Barrientos et al., 2010; Suzuki et al., 2016). Several stress metabolites utilize chorismate as a substrate; although, many of these are typically associated with plant hormone production. We provide evidence in the proteome of UWO241 that a biosynthetic enzyme of the tryptophan pathway, indole-3-glycerol phosphate synthase (IGPS, EC 4.1.1.48) is highly expressed under high salt (Supplemental Table S2). This enzyme is the fourth step in the biosynthesis pathway of L-tryptophan (L-Trp) from chorismite (Fig. 9B). Therefore, it is possible that a product of the shikimate pathway in HS grown UWO241 is the aromatic amino acid L-Trp. However, the metabolome data showed that L-Trp levels were reduced in the HS-grown cultures (Supplemental Table S5). L-Trp is also a major substrate for production of the phytohormone, indole-3 acetic acid (IAA), and the product of IGPS, indol-3-glycerol phosphate, is a branch point between L-Trp synthesis and a L-Trp independent IAA synthesis pathway (Ouyang et al., 2000). IAA and several other phytohomormones have been detected in a few cyanobacteria and algal species; however, their putative function is largely based on exogenously added plant phytohormones to algal cultures (Lu and Xu, 2015). Exogenously added IAA stimulates carbon fixation and growth and enhances stress tolerance in algae (Lu and Xu, 2015). Last, IAA production increases Ca^2+^ levels in plants during acclimation to abiotic stress (Vanneste and Friml, 2013). Ca^2+^ signaling has been linked to both CEF and assembly of PSI supercomplexes (Terashima et al., 2012). Indeed, the Calcium sensing receptor (CAS) was an abundant protein associated with the UWO241-SC (Table 1). More work will be needed to ascertain whether IAA and Ca^2+^ play roles in CEF and assembly of the UWO241-SC.

Despite more than 2 decades of study, the enigmatic UWO241 still has secrets to share. Recent papers have added to the breadth of knowledge on the photosynthetic apparatus, including a cold-adapted ferredoxin isoform (Cvetkovska et al., 2018) and a thylakoid kinase exhibiting temperature-dependent phosphorylation patterns (Szyska-Mroz et al., 2019). Here we extend our understanding of long-term photosynthetic adaptation to permanent cold and hypersalinity by proposing a model for sustained PSI-CEF that supports a robust CBB pathway and a regular growth rate (Fig. 9). Under permanent environmental stress, CEF supplies constitutive photoprotection of PSI and PSII while also producing extra ATP for downstream metabolism (Fig. 9A). The restructured photosynthetic apparatus is accompanied by major rewiring of central metabolism to provide a strong carbon fixation potential which is used in part to produce stored carbon and secondary metabolites (Fig. 9B). Algae adapted to multiple stressors such as low temperatures combined with high salinity are robust fixers of CO_2_, providing new genetic targets for improving crop stress resistance and previously unconsidered sources of natural carbon sinks.

## MATERIALS AND METHODS

### Culture conditions, growth physiology

*Chlamydomonas* sp. UWO241 (UWO241; CCMP1619) was grown in either Bold’s Basal Media (BBM, 0.43 mM NaCl) or BBM supplemented with 700 mM NaCl. Based on earlier studies (Morgan et al. 1998), UWO241 cultures were grown under a temperature/irradiance regime of 8°C/50 photons μmol m^-2^s^-1^. *C.reinhardtii* UTEX 90 was grown in BBM at 20°C and 100 μmol photons m^-2^s^-1^. All cultures were grown in 250 ml glass pyrex tubes in temperature regulated aquaria under a 24 hour light cycle and were continuously aerated with sterile air supplied by aquarium pumps (Morgan-Kiss et al., 2008). Growth was monitored daily by optical density at wavelength of 750 nm. Maximum growth rates were calculated using natural log transformation of the optical density values during the exponential phase. Three biological replicates were performed and all subsequent experiments were conducted on log-phase cultures.

Oxygen evolution was measured at 8 °C with Chlorolab 2 (Hansatech, UK) based on a Clark-type oxygen electrode, following the method described in Jeong et al. (2017) with some modifications. A 2 mL of cell supplemented with 20 μL of 0.5 M NaHCO_3_ was incubated in the dark for 10 min to drain electrons from electron transport chain. The rate of oxygen exchange was measured at increasing light intensities (50, 100, 200, 400 and 600 μmol photons m^− 2^ s^− 1^). Each light lasted 5min and the rates of oxygen evolution at each light intensity step were recorded for 1 min before the end of each light phase. Each light was followed by 2min dark to get the respiration rates.

### Low temperature (77K) fluorescence spectra

Low temperature Chl a fluorescence emission spectra of whole cells and isolated Chl-protein complexes were measured using Luminescence Spectrometer LS50B (Perkin Elmer, USA) as described in Morgan et al. (2008) at 436 nm and 5 (isolate complexes) or 8 nm (whole cells) slit widths. Prior to the measurement, cultures were dark adapted for 10 mins. Decompositional analysis was performed using a non-linear least squares algorithm using Microcal OriginPro Version 8.5.1 (Microcal Origin Northampton, MA). The fitting parameters for the Gaussian components (position, area and full width half-maximum, FWHM) were free running parameters.

### P700 oxidation-reduction and cyclic electron flow

Far red light induced photooxidation of P700 was used to determine rates of CEF as described by Morgan-Kiss et al. (2002b). A volume of exponential phase cultures representing 25 μg Chl a was dark adapted for 10 min and then filtered onto 25 mm GF/C filters (Whatman). Filters were measured on the Dual-PAM 100 instrument using the leaf attachment. The proportion of photooxidizable P700 was determined by monitoring absorbance changes at 820 nm and expressed as the parameter (ΔA_820_/A_820_). The signal was balanced and the measuring light switched on. Far red (FR) light (λmax=715 nm, 10 Wm^−2^, Scott filter RG 715) was then switched on to oxidize P700. After steady-state oxidation levels were reached, the FR light was switched off to re-reduce P700. The half time for the reduction of P700^+^ to P700 (*t*½^red^) was calculated after the FR light was turned off as an estimate of relative rates of PSI-driven CEF (Ivanov et al., 1998). The re-reduction time for P700 was calculated using Microcal™ Origin™ software (Microcal Software Inc., Northampton, MA, USA).

### In vivo spectroscopy measurements

Saturation-pulse chlorophyll fluorescence yield changes and dark interval relaxation kinetics (DIRK) of ECS were measured at 8 °C with the IDEA spectrophotometer as described previously with some modifications (Sacksteder and Kramer, 2000; Zhang et al., 2009). A 2.5 mL of cell supplemented with 25 μL of 0.5 M NaHCO_3_ was pre-incubated in the dark for 10 min and followed by 10 min illumination of far-red light. The chlorophyll fluorescence and ECS were measured with the cells acclimated for 5 min in various actinic light intensities provided by red LEDs. The PSII operating efficiency (Φ_PSII_) was calculated as F_q_’/F_m_’, NPQ as (F_m_-F_m_’)/F_m_’. The linear electron transport (LEF) was calculated from following equation: LEF = *A* × (*fraction_PSII_*) × *I* × Φ_PSII_, where *I* is the light intensity, *A* is the absorptivity of the sample, which is generally assumed to be 0.84 and *fraction_PSII_* is the fraction of absorbed light stimulating PSII (Baker, 2008). The *fraction_PSII_* of UWO241 grown in low-salt and high salt, measured by 77K fluorescence spectra, were 0.709 and 0.746, respectively. The total amplitude of the ECS signal (ECS_t_) was used to estimate the proton motive force (*pmf*). The aggregate conductivity of the thylakoid membrane to protons (*g*_H+_) was estimated from the inverse of lifetime of the rapid decay of ECS (τ_ECS_) (Baker et al., 2007). All ECS signals were normalized to the rapid rise in ECS induced by a single turnover flash to account for changes in pigmentation (Livingston et al. 2010).

### Thylakoid isolation

Thylakoids were isolated according to Morgan-Kiss et al. (1998). Mid-log phase cultures were collected by centrifugation at 2500g for 5 min at 4°C. All buffers were kept ice-cold and contained 1 mM Pefabloc Sc (Sigma, USA) and 20 mM NaF. The pellet was resuspended in grinding buffer (0.3 M sorbitol, 10 mM NaCl, 5 mM MgCl_2_, 5 mM MgCl_2_, 1 mM benzamidine, 1mM amino-caproic acid). The cells were disrupted using chilled French press at 10,000 lb/in^2^ twice, and then and centrifuged at 23,700g for 30 min. The thylakoid pellet was resuspended in wash buffer (50 mM Tricine-NaOH [pH 7.8], 10 mM NaCl, 5 mM MgCl_2_) and centrifuged at 13,300xg for 20 min. The pellet was resuspended in storage buffer (0.3 M sorbitol, 10% glycerol, 50 mM Tricine-NaOH [pH 7.8], 10 mM NaCl and 5 mM MgCl_2_) and stored at −80°C until analysis.

### SDS-PAGE and Immunoblotting

SDS-PAGE was performed using Bio-Rad Mini-Protean system and 12% Urea-SDS gel (Laemmli, 1970). Thylakoid membranes were denatured using 50 mM DTT and incubated at 70°C for 5 min. Samples were loaded on equal protein basis (10 μg total protein). Proteins were transferred to nitrocellulose membrane using cold-wet transfer at 100 V for 2.5 hours. The membrane was blocked with TBST (Tris Buffer Saline Tween) buffer with 5% milk (Carnation). A primary antibody against PsaA (Cat No. AS06-172; Agrisera, Sweden) was used at 1:1000 dilution to probe for major reaction center protein of PSI. Membranes were then exposed to Protein A conjugated to horseradish peroxidase and blots were detected with ECL Select™ Western Blotting Detection Reagent (Amersham).

### Supercomplex isolation

Sucrose step density centrifugation was used to isolate supercomplexes from exponentially grown cultures according to Szyszka-Mroz et al. (2015) with some modifications. Every step was performed in darkness and on ice. All buffers contained phosphatase (20 mM NaF) and protease (1 mM Pefabloc SC) inhibitor. Cells were collected by centrifugation and the pellet was washed twice in Buffer 1 (0.3 M Sucrose, 25 mM Hepes-KOH [pH 7.5], 1mM MgCl_2_). Cells were disrupted using French press, as described above and broken cells were spun down at 50,000g for 30 min. The pellet was resuspended in Buffer 2 (0.3 M Sucrose, 5 mM Hepes-KOH [pH 7.5], 10 mM EDTA) and centrifuged at 50,000g for 30 min. The thylakoid pellet was resuspended gently in Buffer 3 (1.8 M Sucrose, 5mM Hepes-KOH [pH 7.5], 10 mM EDTA) and transferred to Ultra-clear tube (Catalogue No., 344060, Beckman Coulter, USA). The thylakoid prep was overlayed with Buffer 4 (1.3 M Sucrose, 5mM Hepes-KOH [pH 7.5], 10 mM EDTA) followed by Buffer 5 (0.5 M Sucrose, 5mM Hepes-KOH [pH 7.5]). This sucrose step gradient was ultra-centrifuged at 288,000g for 1 hour at 4°C using Sw40Ti rotor (Beckman coulter, USA). Purified thylakoids were collected and diluted (3-fold) in Buffer 6 (5 mM Hepes-KOH [pH 7.5], 10 mM EDTA) and centrifuged at 50,000xg to pellet the membrane. Linear sucrose gradients were made using freeze thaw method with Buffer 7a (1.3 M Sucrose, 5 mM Hepes-KOH [pH 7.5], 0.05% α-DDM) and Buffer 7b (0.1 M Sucrose, 5 mM Hepes-KOH [pH 7.5], 0.05% α-DDM). Briefly, two dilutions of Buffers 7a and 7b were made, Buffer 7-1 (2x Buffer 7a + 1x Buffer 7b) and Buffer 7-2 (1x Buffer 7 a + 2x Buffer 7b). To make the gradient, first 3 ml of Buffer 7a was poured into 12 ml ultra clear tubes followed by flash freezing in liquid nitrogen. Next, Buffer 7-1 was poured on top, followed by flash freezing. This was repeated for Buffer 7-2 and Buffer 7b respectively. The frozen gradients were kept at 4°C overnight to thaw. For supercomplex isolation, thylakoid membranes (0.4 mg Chl) were resuspended in 1% n-dodecyl-alpha-maltoside (α-DDM) (Catalogue number D99020, Glycon Biochemicals, Germany) and incubated on ice in the dark for 25 min. Membranes were spun down to remove insoluble material and loaded onto a linear sucrose gradient described above(0.1 – 1.3 M sucrose) containing 0.05% α-DDM. Gradients were centrifuged at 288,000g for 21 hours at 4°C using SW40Ti rotor (Beckman Coulter, USA). Protein complexes were extracted using a 21-gauge needle.

### Sample preparation for proteomics

Whole cell proteins were extracted as described previously (Valledor and Weckwerth, 2014). Mid-log phase cells were collected by centrifugation at 2500g for 5 min (50 mg wet weight). The cell pellets were resuspended in an extraction buffer containing 100 mM Tris-HCl (pH 8.0), 10% (v/v) glycerol, 2 mM Pefabloc Sc, 10 mM DTT, and 1.2 % (v/v) plant protease inhibitor cocktail (Cat. No. P9599, Sigma). Samples were transferred to 2 mL screw cap tubes containing 25 mg of zirconia beads (Cat. No. A6758, Biorad) and homogenized 3 times for 45 seconds in a BeadBeater (BioSpec). 20% SDS solution was added to the tubes and samples were incubated for 5 min at 95°C. The denatured proteins were centrifuged at 12,000g to pellet any insoluble material. Protein pellets were resuspended in 1.5 ml tris buffer (50mM Tris-HCl, pH 8.0) containing 0.02% n-dodecyl-beta-maltoside (Glycon Biochemicals, Germany) and supplemented with 1X Halt™ protease and phosphatase inhibitor cocktail (Thermo-Scientific, Rockford, IL). After the protein extraction, the sample preparation for proteomics were conducted following our previously published method (Wang et al., 2016). Specifically, 100 μg of total protein were treated with 8 M Urea/5 mM DTT for 1 hour at 37°C, followed by alkylation with 15 mM iodoacetamide in dark for 30 minutes at room temperature. Samples were then diluted four folds with 50 mM Tris-HCl buffer and digested using Mass-spectrometry Grade Trypsin Gold (Promega, Madison, WI) at 1:100 w/w concentration for 16.5 hours at 37°C. The digested samples were cleaned using Sep-Pak C18 plus desalting columns (Waters Corporation, Milford, MA).

### Proteomic analyses by liquid chromatography-tandem mass spectrometry (LC-MS/MS)

The whole cell proteomics were conducted using the Multidimensional Protein Identification Technology (MudPIT) based shotgun proteomics by loading digested peptides onto a biphasic strong cation exchange/reversed phase capillary column. The two dimensional (2D)-LC-MS/MS was conducted on an LTQ ion trap mass spectrometer (Thermo Finnegan, San Jose, CA) operated in the data-dependent acquisition mode. The full mass spectra were recorded at 300-1700 m/z, and the 5 most abundant peaks of each scan were selected for MS/MS analysis. The MS/MS raw data was analyzed by first converting into MS2 files, followed by database search using ProLuCID (Xu et al., 2006). The UWO241 protein database was generated based on our transcriptomics data supplemented with 37 common contaminants, and their reversed sequences as quality control system to restrain false positive discovery to 0.05. Differentially expressed proteins were analyzed using PatternLab for Proteomics (Carvalho et al., 2008). The proteomics results have been deposited to the MassIVE repository with the identifier MSV000084382.

For identifying protein components in the supercomplex, the complex was harvested and 30 μg of total protein was processed similarly as described above to get the digested peptides. Different from the whole cell proteomics, the processed the peptides were directly loaded onto a capillary C18 column without fractionation, and further analyzed in a Thermo LTQ Orbitrap XL mass spectrometer. The full mass spectra were recorded in the range of 350-1800 m/z with the resolution of 30,000. The top 12 peaks of each scan were selected for MS/MS analysis. The data analysis was conducted similarly as described above.

### Identification and analysis of Lhca and BEST protein homologues

Homologues of Lhca and a bestrophin-like protein were identified from a transcriptome previously generated from UWO241 (Raymond et al., 2009; NCBI BioProject No. PRJNA575885). Plastid-targeting signal and transmembrane domains for the putative BEST protein were predicted by ChloroP analysis (http://www.cbs.dtu.dk/services/ChloroP/) and TMHMM software (http://www.cbs.dtu.dk/services/TMHMM-2.0/). The tertiary structure of the C. sp. UWO241 BEST protein was modeled on the high resolution crystal structure of *Klebsiella pneumoniae* Best complex (Yang et al., 2014) in Swiss-Model (https://swissmodel.expasy.org/).

### Gas Chromatography - Mass Spectrometry

For determination of the primary metabolome UWO241 were grown in four biological replicate cultures as described above. Algal cells were harvested by centrifugation (6,000g, 5 min, 4°C) and washed once with fresh media. The supernatant was decanted, and the algal cells were flash frozen in liquid nitrogen and stored at −80°C. The metabolite extraction protocol was adapted from (Fiehn et al., 2008). In brief, metabolites were extracted from 20 mg of frozen tissue in 1 ml cold extraction buffer (methanol: chlroform: dH_2_O; 5:2:2). The samples were homogenized using glass beads (500 μm i.d.) in a Geno/Grinder 2010 instrument (SpexSamplePrep, Metuchen, NJ, USA), followed by centrifugation (14,000g, 2 min, 4°C). Samples were further processed and derivatized for GC-TOF mass spectrometry as described (Lee and Fiehn, 2008). GC-MS measurements were carried out on an Agilent 6890 gas chromatograph (Agilent, Santa Clara, CA, USA), controlled by a Leco ChromaTOF software v 2.32 (Leco, St. Joseph, MI, USA). Separation was performed on a Rtx-5Sil MS column (30m x 0.25mm x 0.25μm) with an additional 10 m empty guard column (Restek, Bellefonte, PA, USA) using helium as a carrier (1 ml/min flow rate). The oven temperature was held constant at 50°C for 1 min, the ramped at 20°/min to 330°C at which it was held constant for 5 min. A Leco Pegasus IV mass spectrometer (Leco, St. Joseph, MI, USA) was operated in electron impact (EI) mode at −70 eV ionization energy with unit mass resolution at 17 spectra/s with a scan range of 80-500 Da. The transfer line temperature between gas chromatograph and mass spectrometer was set to 280°C. Ionization Electron impact ionization at 70V was employed with an ion source temperature of 250°C. Mass spectra were processed using BinBase, an application system for deconvoluting and annotating mass spectral data, and analyzed as described in (Fiehn et al., 2005). Metabolites were identified based on their mass spectral characteristics and GC retention times, by comparison with compounds in a plant and algae reference library (West Coast Metabolomics Center, UC Davis, CA, USA). Peak heights for the quantification ion at the specific retention index corresponding to each metabolite were normalized by the sum of peak heights in the sample. Normalized data were processed by cube root transformation followed by range scaling (van den Berg et al., 2006). Statistical analyses were performed by the Metaboanalyst 4.0 software suite (Chong et al., 2018), and included principal component analysis (PCA), t-test, heatmap and clustering analysis using Ward’s linkage for clustering and Pearson’s correlation as a measure of dissimilarity.

## ACKNOWLEDGEMENTS

The authors thank the staff (Dr. Andor Kiss and Ms. Xiaoyun Deng) of the Center for Bioinformatics & Functional Genomics (CBFG) at Miami University for instrumentation and computational support. We also would like to thank people from the Kramer Laboratory (Drs. David Kramer, Jeffrey Cruz, Robert Zegarac, Ben Lucker) for their suggestions to setup and optimize the IDEAspec for spectroscopic measurements in UWO241. Funding was provided by NSF Grant 1637708 (RMK, IK), DOE Grant DE-SC0019138 (RMK, IK, XW), DOE grant DE-SC0019464 (JJ, WM, RZ) and Donald Danforth Plant Science Center startup fund (RZ).

